# Fast Numerical Optimization for Genome Sequencing Data in Population Biobanks

**DOI:** 10.1101/2021.02.14.431030

**Authors:** Ruilin Li, Christopher Chang, Yosuke Tanigawa, Balasubramanian Narasimhan, Trevor Hastie, Robert Tibshirani, Manuel A. Rivas

## Abstract

We develop two efficient solvers for optimization problems arising from large-scale regularized regressions on millions of genetic variants sequenced from hundreds of thousands of individuals. These genetic variants are encoded by the values in the set {0, 1, 2, `NA`}. We take advantage of this fact and use two bits to represent each entry in a genetic matrix, which reduces memory requirement by a factor of 32 compared to a double precision floating point representation. Using this representation, we implemented an iteratively reweighted least square algorithm to solve Lasso regressions on genetic matrices, which we name `snpnet-2.0`. When the dataset contains many rare variants, the predictors can be encoded in a sparse matrix. We utilize the sparsity in the predictor matrix to further reduce memory requirement and computational speed. Our sparse genetic matrix implementation uses both the compact 2-bit representation and a simplified version of compressed sparse block format so that matrix-vector multiplications can be effectively parallelized on multiple CPU cores. To demonstrate the effectiveness of this representation, we implement an accelerated proximal gradient method to solve group Lasso on these sparse genetic matrices. This solver is named `sparse-snpnet`, and will also be included as part of `snpnet` R package. Our implementation is able to solve group Lasso problems on sparse genetic matrices with more than 1, 000, 000 columns and almost 100, 000 rows within 10 minutes and using less than 32GB of memory.

## 1 Introduction

Constantly growing biobanks have provided scientists and researchers with unprecedented opportunities to understand the genetics of human phenotypes. One component is to predict phenotypes of an individual using genetic data. However, datasets of increasing size also pose computational challenges for this task. On the statistics side, genetic datasets are usually high-dimensional, meaning the number of genetic variants is larger than the number of sequenced individuals. High dimensional statistics have been studied for more than two decades with well understood solutions. One such solution is to “bet on sparsity”: the assumption that only a small subset of variables are associated with the response. The sparsity assumption is usually embodied through an objective function that encourages sparsity in the solution. Well known examples include the Lasso and the group Lasso. On the computation side, a statistical estimator that describes the relationship between the genetic variants and the response of interest are often obtained by optimizing an objective function involving the genetic matrix. While off-the-shelf solvers may exist for these optimization problems, they are usually not optimal for genetics data. First, these general purpose solvers require loading a floating point predictor matrix in memory before optimization can be done. This can demand a very large amount of memory for biobank scale data. For example, loading a matrix with 200, 000 rows and 1, 000, 000 columns as double precision floating point numbers takes 1.6 terabytes, much larger than the RAM size of most machines. In particular, they do not exploit the fact that genetic variants can take on only four possible values. Secondly, many of these solvers do not fully utilize modern hardware features such as multi-core processors, which leaves lots of performance on the table. Thirdly, a large number of variants in exome and whole genome sequencing data are rare variants. If a variant is encoded as the number of copies of the minor allele, then the corresponding genetic matrix is sparse. In the UK Biobank’s exome sequencing data (Szustakowski et al. 2020), more than 99% of the variants in the targeted regions have minor allele frequency less than 1%. As a result, more than 98% of the entries of the corresponding genetic matrix are zero. The sparsity in the predictor matrix can potentially be exploited to improve both memory requirements and computational speed.

The main result of this work is an extremely efficient regularized regression solver for problems with sparse genetic predictors, named `sparse-snpnet`. The main features of this solver are the following:

1. A compact, two bits representation of genetic variants based on PLINK2’s (Chang et al. 2015) pgen files.
2. Good scalability to multi-core processors.
3. A simplified version of the compressed sparse block format so that arithmetic operations on the genetic matrices are more amenable to parallelism.

In addition, we provide an extension to the popular R package `glmnet` (Friedman et al. 2010, Simon et al. 2011) specifically for Lasso problems involving genetic matrices. This extension exploits the compact representation and is multi-threaded, but does not assume sparsity of the input genetic matrix. We incorporate this solver to the screening framework (Qian et al. 2020) in `snpnet` and name it `snpnet-2.0`. Both solvers are implemented in C++ and wrapped as part of the R package `snpnet`, which is available at https://github.com/rivas-lab/snpnet/tree/compact. We refer the readers to section 5 for comparisons between these two methods.

## 2 Results

### 2.1 Optimization Algorithm

We focus on regularized regression problems whose objective functions are in the following form:

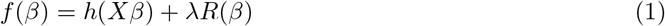

where *X* ∈ {0, 1, 2, NA}^*n*×*d*^ is a genetic matrix, β ∈ ℝ^*d*^is the parameter vector, *h*: ℝ^*n*^↦ ℝ is usually the negative log-likelihood function of a generalized linear model (Hastie & Tibshirani 1986), and is always assumed to be smooth and convex. We have omitted the dependence of *h* on the response vector to simplify the notation. *R*: ℝ^*d*^↦ ℝ_+_ is a regularization function, and λ ∈ R_+_ represents the strength of regularization. Here are some examples of *h*:

1. Linear regression: 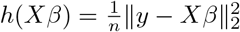 for a response vector *y* ∈ ℝ^*n*^.
2. Logistic regression: write 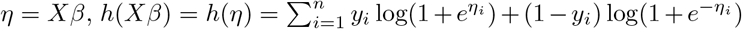 for a binary response *y* ∈ {0, 1}^*n*^.
3. Cox regression (Cox 1972): Write 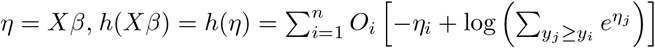 for a survival time vector 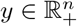 and an event indicator *O* ∈ {0, 1}^*n*^.

The regularization function is usually a seminorm but not always. Some examples are:

1. Lasso (Tibshirani 1996): 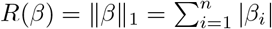.
2. Elastic net (Zou & Hastie 2005): 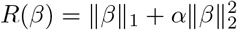 for some *α* > 0.
3. Group Lasso (Yuan & Lin 2006): *R*(*β*) = ∑_*g*∈𝒢_ ∥*β*_*g*_∥_2_, where *g*∈𝒢, *g* ⊆ {1,2,…, *d*} represents a subset of variables.

To minimize (1), we apply an accelerated proximal gradient descent algorithm (Nesterov 1983, Daubechies et al. 2004, Beck & Teboulle 2009) with backtracking line search to determine the step size. This algorithm has fast convergence rate, essentially no tuning parameter, and is particularly suitable for the simple regularization functions that we use. In short, this algorithm alternates between a gradient descent step that decreases the value of *h*(*Xβ*), and a proximal step that ensures that the regularization term is not too large. Note that the gradient here refers to the gradient of *h*(*Xβ*) with respect to *β*. The regularization function is usually not differentiable at 0. The proximal operator is defined as:

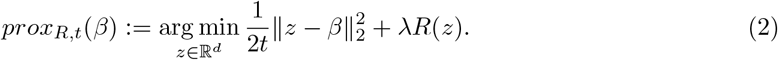

When the regularization function is one of the examples above, the corresponding proximal operator have explicit expression. We summarize this process in the pseudo-code in algorithm 1.

We observe that in this algorithm, the only operations that involve the predictor matrix *X* are matrix-vector multiplications *Xβ* and *X*^*T*^ *r*, where *r* = *∇h*(*Xβ*) ∈ ℝ^*d*^. When *X* is dense, these two operations are also the most computationally intensive ones in this algorithm, having complexity 𝒪(*nd*), whereas all other operations are either 𝒪 (*n*) or 𝒪 (*d*). This, together with the need to reduce the amount of memory required to load *X*, motivate a more compact and efficient representation of the genetic predictor matrix *X*.

### 2.2 Sparse Genotype Matrix Representation

In this section we describe the format we use to represent sparse genetic matrices. First of all, we pack each entries in the matrix to two bits. 0, 1, 2 and `NA` are represented by 00, 01, 10 and 11, respectively. The compressed sparse column (CSC) format is a popular way to store a sparse matrix. The PLINK 2.0 library (Chang et al. 2015) provides functions that make loading a genetic matrix into this format straightforward. Under CSC, a matrix with *n* rows, *d* columns and *nnz* non-zero entries are represented by three arrays:

1. A column pointer array `col ptr` of size *d* + 1.
2. A row index array `row idx` of size *nnz*.
3. A value array `val` of size *nnz*.

For each column *j* ∈ {1, 2, *…, d*}, the non-zero entries in that column are stored from the `col ptr[j]`th (inclusive) entry to the `(col ptr[j+1] - 1)`th entry of `row idx` and `val`, where `row idx` stores the row index of the non-zero entry and `val` stores the non-zero value. Figure 1 provides an illustration of a sparse genetic matrix under CSC format.

**Figure 1.**
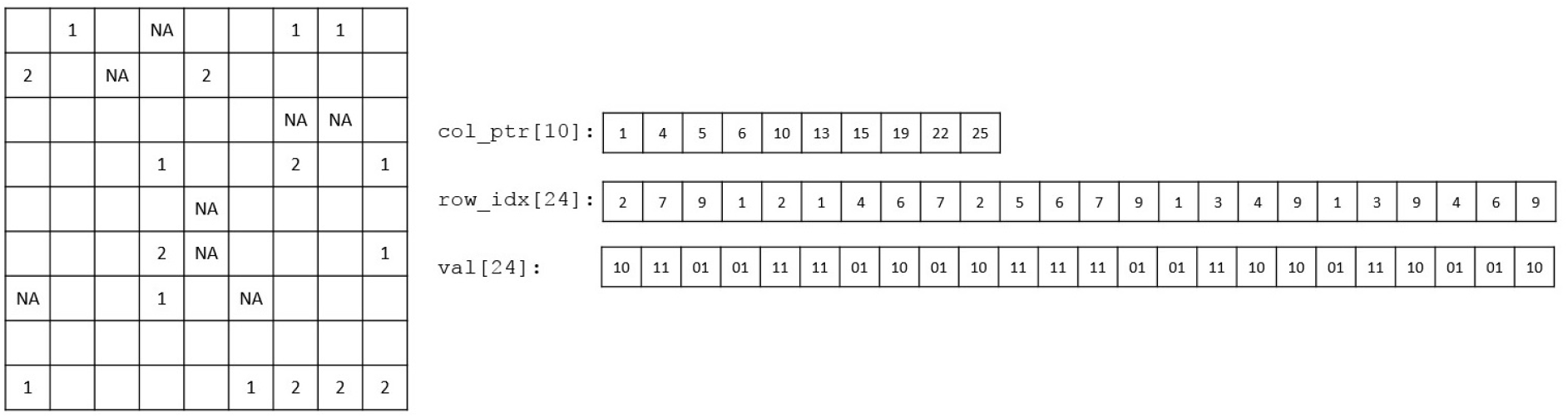
A sparse genetic matrix represented in the compressed sparse column format. The zero entries on the left are omitted.

When a sparse matrix is stored in CSC format, accessing a particular column is simple. As a result, one can trivially parallellize the computation of *X*^*T*^ *r*. For example, thread *j* can compute the inner product of the *j*th column of *X* and *r* and write to the *j*th entry of the output without interfering with other threads. However, same thing can’t be said about *Xβ*. We can’t directly access a row of *X* stored in CSC format, so there is no easy way to make each thread compute the inner product between *β* and a row of *X*. Another way is to, say, have thread *j* add *β*_*j*_ times the *j*th column of *X* to the output, but doing this in parallel leads to data race. Alternatively, one can store *X* in the compressed sparse row format, which makes parallelizing *Xβ* easy but *X*^*T*^ *r* difficult.

Our implementation uses a simplified version of the compressed sparse block (CSB) format proposed in Buluç et al. (2009). In this format, the sparse matrix is partitioned into a grid of smaller, rectangular sub-matrices with same dimensions, which are referred to as blocks. When partitioned to *B* blocks, a matrix with *n* rows, *d* columns, and *nnz* non-zero entries are represented by four arrays:

1. A block pointer array `blk ptr` of size *B* + 1.
2. A row index array `row idx` of size *nnz*.
3. A column index array `col idx` of size *nnz*.
4. A value array `val` of size *nnz*.

In this representation, non-zero entries in a block (as oppose to those in a column in CSC format) are stored contiguously. The row indices, column indices, and values of the non-zero values in a block *b* ∈ {1, 2, …, *B*} are stored in `row idx`, `col idx`, and `val`, starting at the index `blk ptr[b]` and ending at the index `blk ptr[b+1]-1`. In the original CSB paper the non-zero elements in each block has a Z-Morton ordering, while the blocks can have any order. In our simplified version we store the blocks and the non-zero elements within a block in a column major fashion. Figure 2 provides an illustration.

**Figure 2.**
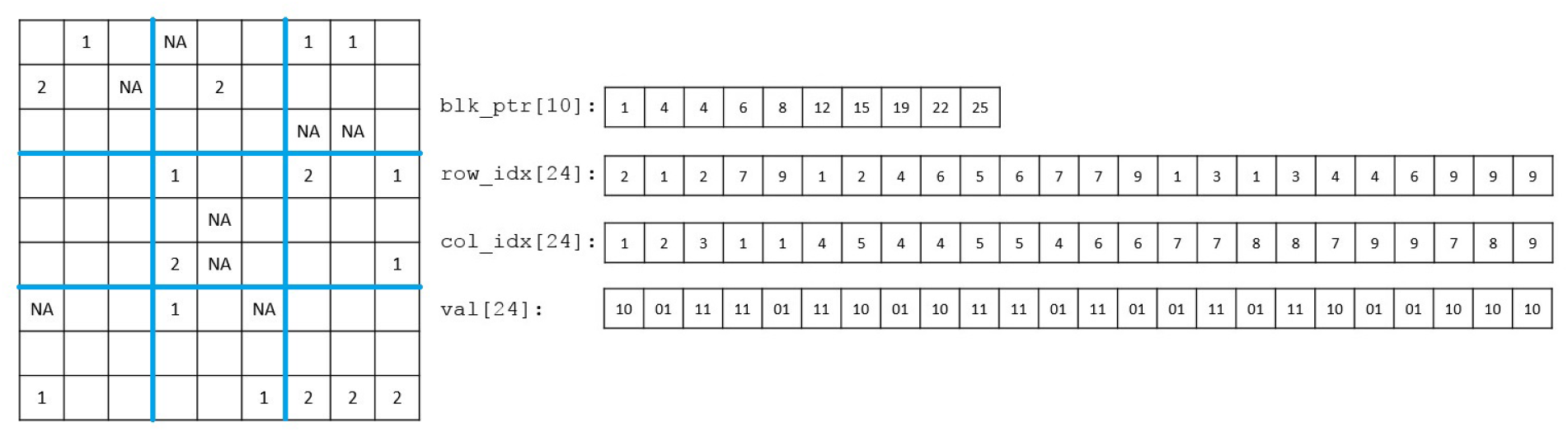
A sparse genetic matrix represented in the compressed sparse block format. The zero entries on the left are omitted. The boundaries of each block are highlighted in blue. The non-zero elements in each block are stored contiguously in an top-to-bottom, left-to-right order. The blocks are also in an top-to-bottom, left-to-right order

Under this representation accessing a block in the sparse matrix is easy. As a result, parallellizing both *Xβ* and *X*^*T*^ *r* are straightforward. For example if *X* is the matrix in figure 2, then to compute *Xβ* we can have thread 1 compute the inner product of the first three rows of *X* and *β*, thread 2 compute the inner product of row 4-6 and *β*, etc. Similarly, to compute *X*^*T*^ *r* thread *b* will compute the inner product between *r* and the columns 3*b −* 2, 3*b −* 1, 3*b* in *X* for *b* ∈ {1, 2, 3}.

Since our implementation uses 2 bits to store a matrix entry and 32 bits to store each index, the genetic CSB format (compared to a dense representation) will only save memory when the matrix is sufficiently sparse (approximately *<* 3% of entries are non-zero). While there are many techniques to reduce the number of bits needed to represent the indices (such as storing the indices relative to the start of the block, differential encoding, bit packing), the current version of our software does not implement these techniques. For variants with high minor allele frequency, we store the corresponding columns in dense format and keep track of their column indices. We also keep an additional array of length *d* to store the mean imputation of the missing values in *X*.

#### Algorithm 1 Accelerated Proximal Gradient Method for (1)

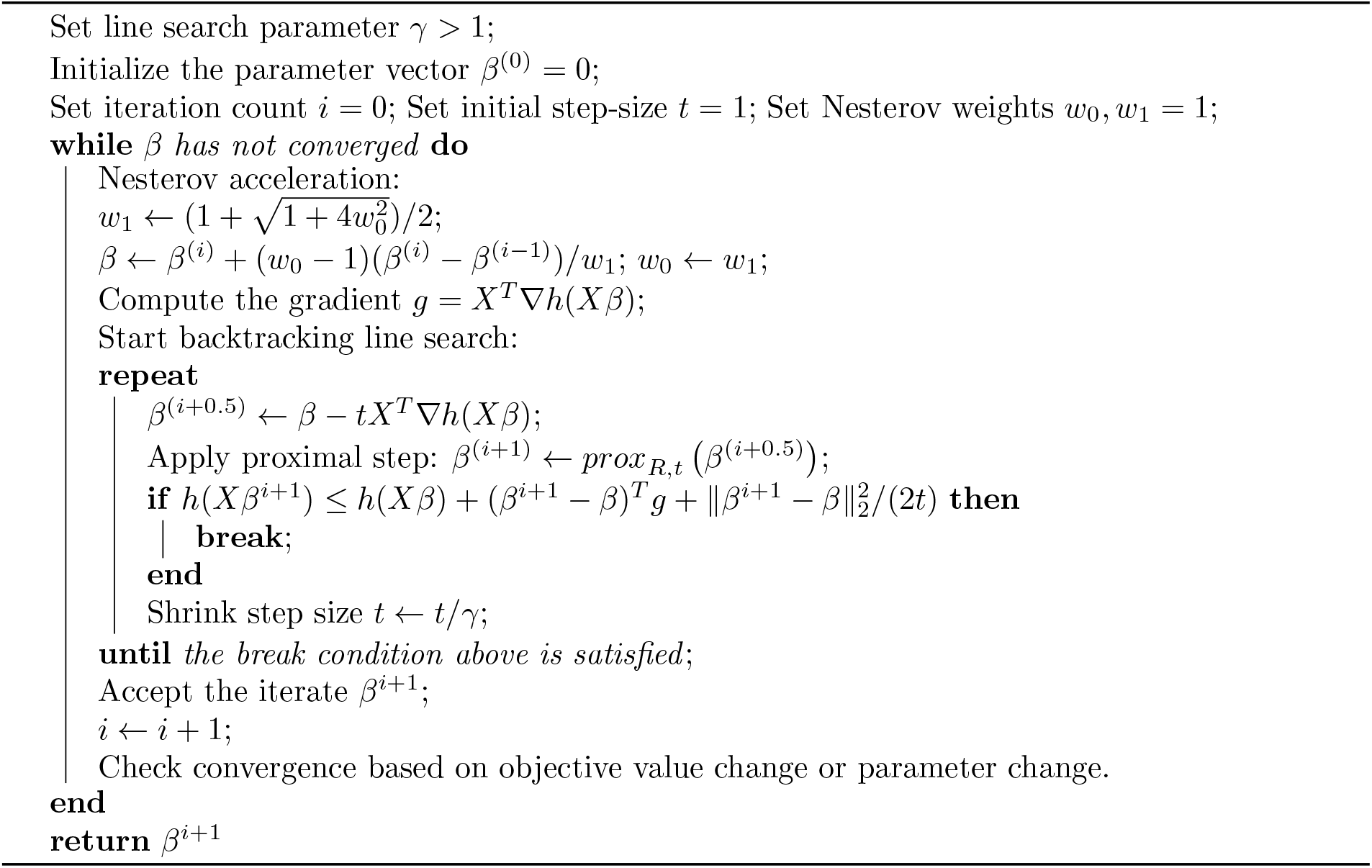

## 3 Benchmarks

### 3.1 Performance on Dense Matrix-Vector Multiplications

In the first benchmark we evaluate the performance improvement when the predictor matrix uses the 2-bit compact representation, but not the sparse format described in the last section. The genetics data are dense and simulated through the `plink2 --dummy` command. The matrix have *n* = 200, 000 rows and *d* = 30, 000 columns with approximately 5% entries `NA`s. In Figure 3 we show the relative speedup of the compact matrix as a function of the number of threads used. The baseline is R’s builtin matrix-vector multiplication functions for double precision matrices (a basic, single threaded BLAS implementation). The numbers reported are based on the median wall time of 10 runs. Figure 3 demonstrates a more than 20-35 folds of speedup over the baseline, and good performance scalability in the number of threads for up to almost 20 threads. Unless otherwise specified, all computational experiments in this paper are done on an Intel Xeon Gold 6258R. For most of our applications 16 out of the 28 CPU cores that comes with this CPU are used.

**Figure 3.**
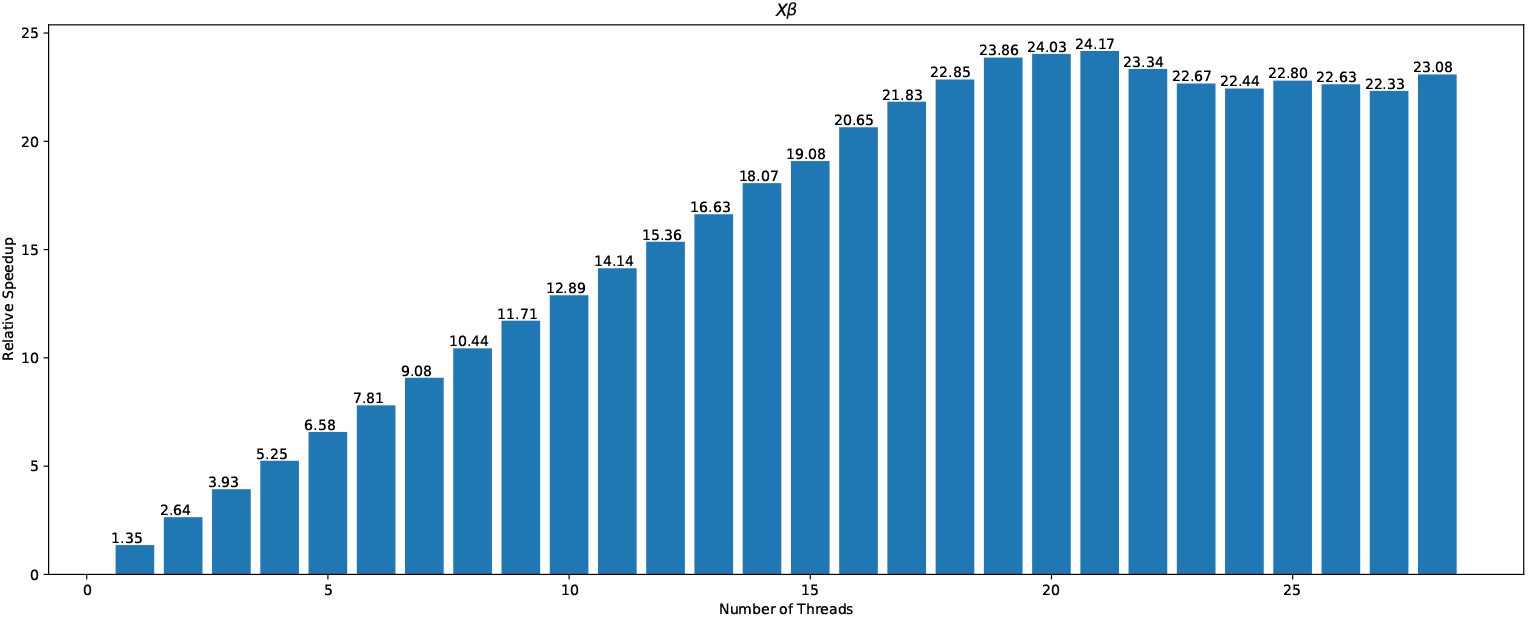
A bar plot demonstrating the relative speedup in computing *Xβ* when the compact representation is used. The baseline is the `R`’s builtin matrix-vector multiplication function for double precision matrices. The horizontal axis is the number of threads. The vertical axis is the ratio of between time spent in the baseline and the time spent with the compact representation. The baseline for *Xβ* is 9.8 seconds.

**Figure 4.**
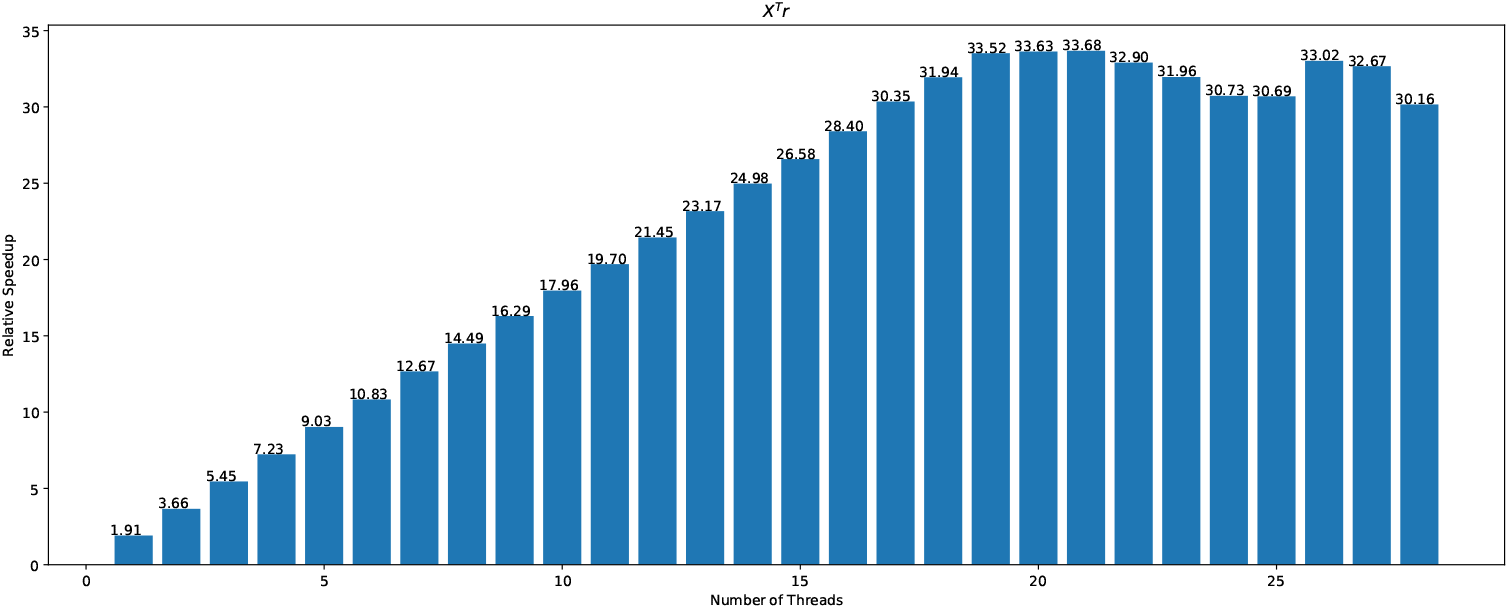
A bar plot demonstrating the relative speedup in computing *X*^*T*^ *r* when the compact representation is used. The baseline is the `R`’s builtin matrix-vector multiplication function for double precision matrices. The horizontal axis is the number of threads. The vertical axis is the ratio of between time spent in the baseline and the time spent with the compact representation. The baseline for *X*^*T*^ *r* is 11.4 seconds.

### 3.2 Performance on Solving Large-scale Lasso Problems

In the second benchmark we compare the performance of the 2-bit compact genetic matrix representation when it is incorporated in the software packages `glmnet` and `snpnet` to solve large-scale Lasso problems where the predictors are mostly Single-nucleotide polymorphisms (SNPs) (with perhaps a few real-valued covariates such as age). As mentioned in the introduction, we call this implementation `snpnet-2.0`. The main performance gain comes from these factors (the readers can refer to Qian et al. (2020), Li et al. (2020) for the definitions of some terms below):

1. `glmnet` fitting is based on the iteratively reweighted least square (IRLS) algorithm, where the main bottleneck is computing inner-products between columns of *X* and a real-valued vector. This is done using the 2-bit compact representation and is multi-threaded in `snpnet-2.0`.
2. `snpnet` uses a screening procedure named the batch screening iterative Lasso (BASIL). At each BASIL iteration a different set of predictors is used to fit a model. Using the compact representation reduces the amount of memory traffic needed.
3. `snpnet-2.0` uses reduced precision floating point numbers (`float32` instead of `float64`) to do KKT checking.
4. Warm start support, as well as more relaxed convergence criteria, for binomial model and Cox model.

Since `snpnet-2.0` also uses more relaxed convergence criteria than the previous version, it is not very fair to just compare the speed of these packages. As a result we provide both the time spent as well as the test set prediction performance. The data used here are a combination of directly genotyped variants (release version 2 of Sudlow et al. (2015)), the imputed allelotypes in human leukocyte antigen allelotypes (Venkataraman et al. 2020), and copy number variations described in Aguirre et al. (2019), resulting in a genotype matrix of 1, 080, 968 variants, as described in Sinnott-Armstrong et al. (2021). The study population consists of 337, 129 unrelated participants of white British ancestry described in DeBoever et al. (2018). We randomly select 70% of the participants as the training set, 10% as the validation set, and 20% as the test set. The results are summarized in table 3. The table shows that `snpnet-2.0` achieves significant speedup over the old version while having very similar test set prediction performance. We note that for standing height, more than 80, 000 variants are selected to fit the model. Since the predictor matrix is duplicated in its fitting process, the old version of `snpnet` requires more than 400GB to successfully finish. On the other hand, 32GB of memory is sufficient for `snpnet-2.0`.

**Table 1:** Speed comparison between the old version of `snpnet` and `snpnet-2.0`. Time is measured in **minutes**. (B) indicates the response is binary, (Q) indicates the response is quantitative, and (S) indicates that the response is a survival time. For binary response, the test metric is the area under the ROC curve (AUC). For quantitative response, the metric is the R-squared. For survival response, the metric is the C-index. ^*^The machine we used for most of the applications here has a dual-socket architecture, each having around 400 GB of local memory. The memory requirements by the old version of `snpnet` for both standing height and BMI exceeds the capacity of the local memory of a single socket in this machine. As a result, we ran these two experiments on an Intel Xeon Gold 6130 (also 16 cores) machine with more memory.

|  | Time (new) | Time (old) | Test metric (new) | Test metric (old) |
| --- | --- | --- | --- | --- |
| High cholesterol (B) | 21.9 | 109.8 | 0.72533 | 0.72531 |
| Asthma (B) | 21.7 | 130.0 | 0.61609 | 0.61608 |
| Standing Height (Q) | 99.9 | 405.8* | 0.71096 | 0.71100 |
| BMI (Q) | 51.5 | 208.3* | 0.11408 | 0.11412 |
| Other hypothyroidism (S) | 13.5 | 71.5 | 0.75194 | 0.75205 |
| Thyrotoxicosis (S) | 3.6 | 10.0 | 0.71020 | 0.71021 |

**Table 2:** The loading and computation time in **seconds** when the genetic matrix is stored in sparse format. This matrix has 200, 643 rows and 7, 462, 671 columns with more than 10 billion non-zero entries. The computation time are the median of 10 runs. Both dense and sparse format compute the matrix-vector multiplication using 16 cores.

| Loading | $X\beta$ | $X^T r$ |
| --- | --- | --- |
| 56.5 | 1.86 | 1.80 |

**Table 3:** A comparison between the two solvers we present in this paper.

|  | `snpnet-2.0` | `sparse-snpnet` |
| --- | --- | --- |
| Algorithm | IRLS | Proximal gradient |
| Use variable screening | Yes | No |
| Easy to extend to other GLMs | Yes | Yes |
| Easy to extend to other regularizations | No | Yes |
| Use 2-bit representation of variants | Yes | Yes |
| Use sparse matrix format | No | Yes |
| Multi-threaded | Yes | Yes |

### 3.3 Performance of the Sparse Format

In the third benchmark we evaluate the performance improvement when the genetic predictors make use of both the 2-bit compact representation, and the sparse representation described in the last section. In this case we use real exome data from the UK Biobank. The raw data has 200, 643 individuals and 17, 777, 950 variants. For this benchmark we only use variants with least 3 individuals having the minor allele and with missing rate at most 10%. The result is a sparse genetic matrix with 200, 643 rows and 7, 462, 671 columns. For our application in the next section the number of variants used to fit models will be smaller since the training set will be a subset of the entire population in this data. On average each column of this matrix has 1399.5 non-zero entries, half of the columns have less than 7 non-zero entries, and 90% of the columns have less than 91 non-zero entries. In our sparse representation we divide this matrix into 16 *×* 16 = 256 blocks, each with dimension 12, 540 by 466, 416 (the size of the blocks is a tuning parameter), except at the boundary the block size could be larger. As we mentioned in the last section, storing dense blocks using our version of the compressed sparse block format is not memory efficient, so if a column has a large number of non-zero entries we store all entries of that column separately. For this particular matrix, 223, 596 variants does not use the sparse representation. In table 2 we present the amount of time to load the matrix and to compute *Xβ, X*^*T*^ *r* using the sparse matrix representations. Again 16 cores are used for the computation. Loading such matrix would take almost 12 terabytes of memory if the entries are stored as double precision floating point numbers.

## 4 Applications to UK Biobank Exome Sequencing Data

In this section we put our method into practice. Specifically, we use the exome data described in the last part of section 3 to fit group-sparse linear models on multiple phenotypes. The method described in this section is implemented in `sparse-snpnet`. In this case the regularization term will be the sum of the 2-norms of the predefined groups. For our applications the groups are defined by the gene symbol of the variants. For example, the objective function for a Gaussian model is:

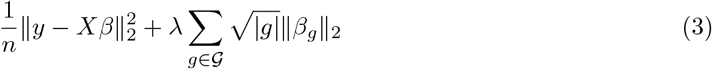

where 𝒢 = *{g*: *g ⊆* {1, 2, *…, d*}} is a collection of indices corresponding to variants with the same gene symbol. |*g*|, the number of element in the group, is part of the regularization term so that groups of same size are penalized by the same degree. *β*_*g*_ ∈ ℝ^|*g*|^ is the sub-vector of *β* corresponding to the indices in *g*. We do not allow overlapping groups, so 𝒢 needs to be a partition of all variables. That is *∪*_*g*∈𝒢_ *g* = {1, 2, *…, d*}, ∑_*g*∈𝒢_ |*g*| = *d*. One can show that the proximal operator for this and regularization function satisfies:

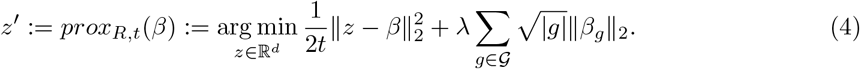

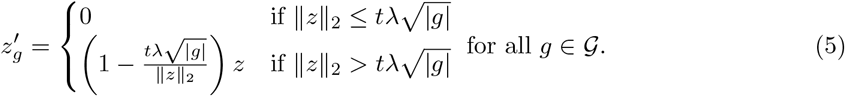

In practice we would like to adjust for covariates such as age, sex, and other demographic information when fitting a regression model. In our application we first fit a unregularized regression model of the response on these covariates and fit the regularized model of the residual on the genetic variants. The number of covariates are usually much smaller compared to the number of individuals, so the first fitting is not computationally or statistically challenging. To be more precise, let *X*_*cov*_ ∈ ℝ^*n×c*^ be the *c* ≥ 0 covariates that we would like to adjust for. Using the same notation in (1). We fit a model in two steps:

1. First, we fit a unregularized model using the covariates:

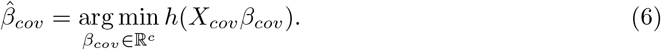

1. Then, we fit the regularized model on the “residuals” using the variants:

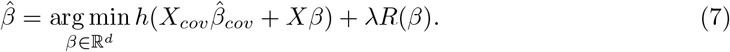

For example, if we write the predicted value of the covariates as 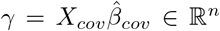, then for a Gaussian model, the objective function of the second step above is:

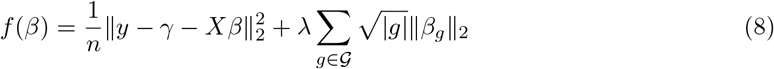

For logistic regression this becomes:

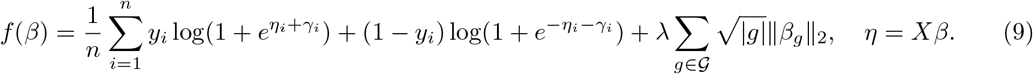

For Cox model the objective function is

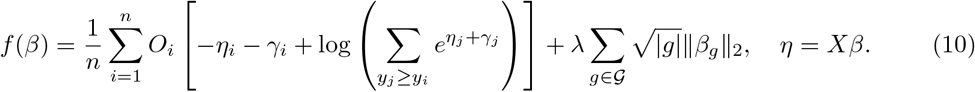

We optimize these objective functions for a decreasing sequence of *λ*s starting from one such that the solution just becomes non-zero. The initial value for the proximal gradient method of the next *λ* are initialized from the solution from the current *λ* (warm start). As alluded in the last section, we randomly assign 70% of the white British individuals in this dataset to the training set, 10% to the validation set, and 20% to the test set. We remove individuals whose phenotype value is missing, keeping variants with at least 3 minor allele count, has an associated gene symbol, has less than 10% of missing value, and the ratio of the missing value and minor allele is less than 10. We further filter out the variants that are not protein truncating or protein altering. Depending on the number of missing values in the phenotype, the number of individuals and variants used for fitting could be different. In all of the examples here the training set has more than 90, 000 individuals and the number of genetic variants used are over 1, 000, 000. The covariates are the sex, age, and 10 principal components of the genetic data described in the second benchmark of section 3.2.

To evaluate the fitted model, we use the R-squared value for quantitative phenotype, the area under the receiver operating characteristic (ROC) curve (AUC) for binary phenotype, and the concordance index (C-index) for time-to-event phenotype. These metrics will be computed on the validation set to determine the optimal regularization parameter *λ* and on the test set to evaluate the model corresponding to the *λ* used. Once the validation metric starts to decrease, we stop the fitting process and do not compute the solutions for smaller *λ* values. Figures 5, 6, 7 illustrate a few Lasso path plots obtained from our implementation. It is worth noting that Alzheimer’s disease has a sharp increase in C-index for *λ* indices 7-10.

**Figure 5.**
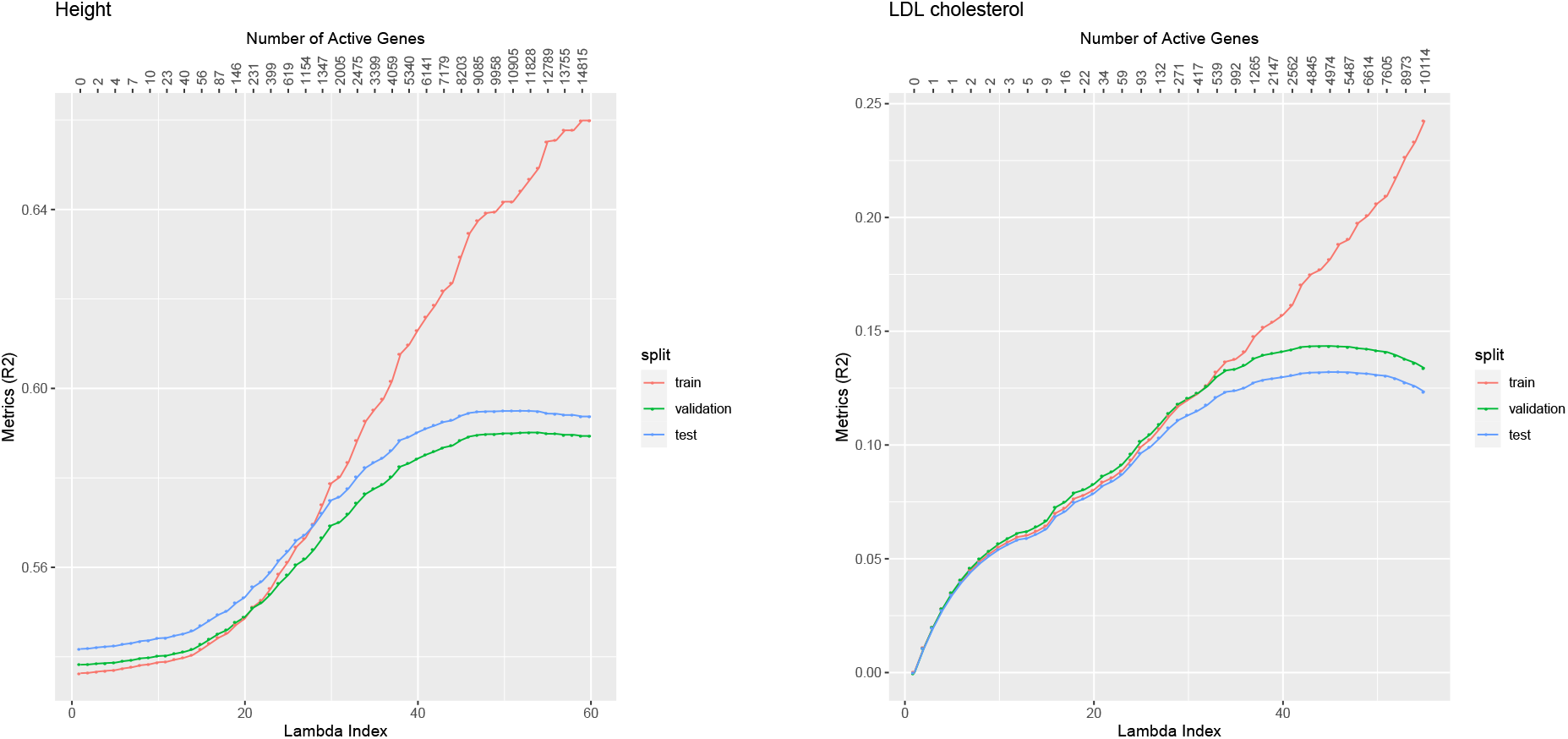
Lasso path plots for the quantitative phenotypes Height and LDL cholesterol. The horizontal axis is the index of the regularization parameter *λ*, the vertical axis are the R-squared of the solution corresponding to each *λ* index. The numbers on the top are the number of genes with non-zero coefficients at the corresponding *λ* indices. The color corresponds to the train, validation and test set. The duration of training these two models are 8.34 minutes and 8.35 minutes respectively.

**Figure 6.**
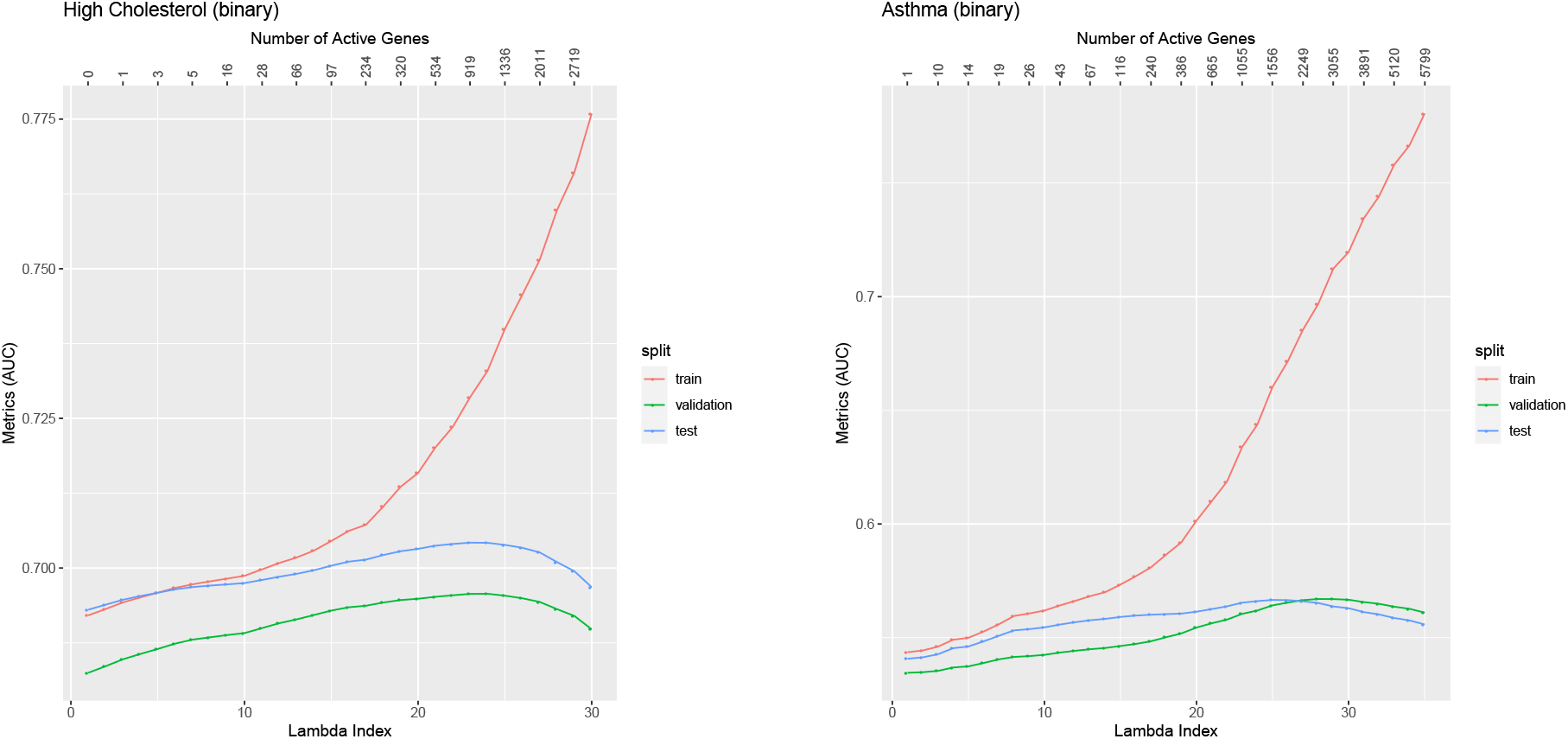
Lasso path plots for the binary phenotypes High Cholesterol and Asthma. The horizontal axis is the index of the regularization parameter *λ*, the vertical axis are the AUC values of the solution corresponding to each *λ* index. The numbers on the top are the number of genes with non-zero coefficients at the corresponding *λ* indices. The color corresponds to the train, validation and test set. The duration of training these two models are 6.88 minutes and 8.27 minutes respectively.

**Figure 7.**
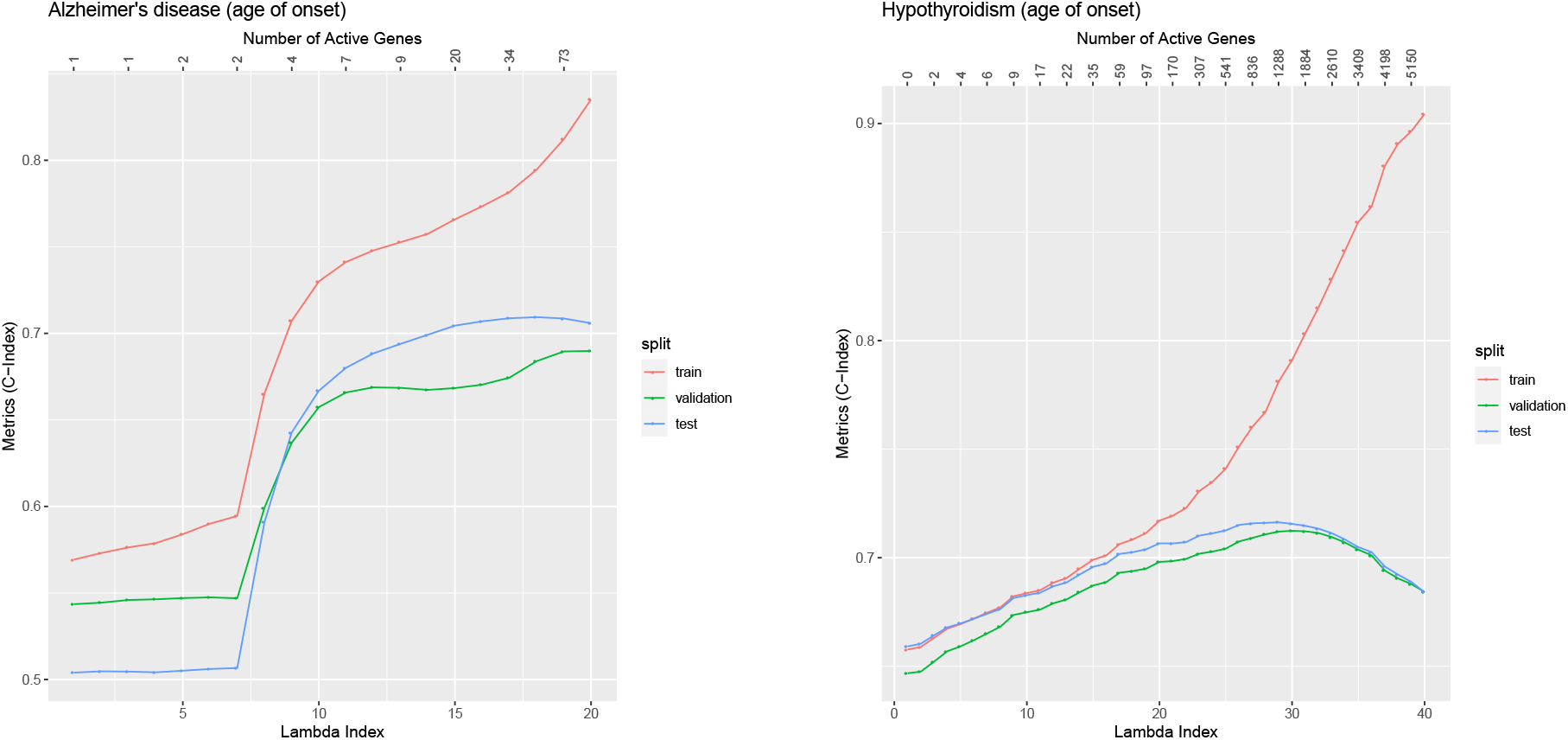
Lasso path plots for the time-to-event phenotypes Alzheimer’s disease and Hypothyroidism. The horizontal axis is the index of the regularization parameter *λ*, the vertical axis are the Cindex of the solution corresponding to each *λ* index. The numbers on the top are the number of genes with non-zero coefficients at the corresponding *λ* indices. The color corresponds to the train, validation and test set. The duration of training these two models are 6.12 minutes and 6.92 minutes respectively.

In terms of computation, unlike in `snpnet` (or the `2.0` version), the optimization in `sparse-snpnet` are all done without variable screening, and the entire training data (in sparse format) is loaded in memory before fitting starts. This eliminates all the I/O operations carried out in the KKT checking step of `snpnet`. The applications in this section successfully finished when we allocate 32GB of memory to these jobs. In addition, while the applications in this paper focus on Gaussian, logistic, and Cox families and group Lasso regularization, our implementation uses several abstraction in C++ so it’s easy to extend to other generalized linear models and regularization functions.

## 5 Discussions

We present two fast and memory efficient solvers for generalized linear models with regularization on large genetic data. Both methods utilize a 2-bit compact representation of genetic variants and are accelerated though multi-threading on CPUs with multiple cores. The first solver implements the iteratively-reweighted least square algorithm in `glmnet`, and its goal is to provide boosted computational and memory performance to the large-scale Lasso solver described in (Qian et al. 2020, Li et al. 2020). The second solver implements an accelerated proximal gradient method that’s able to solve more general regularized regression problems. One important feature of this solver is that it combines a version of compressed sparse block format for sparse matrices and the 2-bit encoding of genetic variants. We summarize the characteristics of these two solvers in table 3. We demonstrate the effectiveness of our methods through several benchmarks and UK Biobank exome data applications. We believe our method will be a useful tool as whole genome sequencing data becomes more common.

## 6 Acknowledgments

Y.T. is supported by a Funai Overseas Scholarship from the Funai Foundation for Information Technology and the Stanford University School of Medicine.

M.A.R. is supported by Stanford University and a National Institute of Health center for Multi and Trans-ethnic Mapping of Mendelian and Complex Diseases grant (5U01 HG009080). This work was supported by National Human Genome Research Institute (NHGRI) of the National Institutes of Health (NIH) under awards R01HG010140. The content is solely the responsibility of the authors and does not necessarily represent the official views of the National Institutes of Health.

R.T was partially supported by NIH grant 5R01 EB001988-16 and NSF grant 19 DMS1208164.

T.H. was partially supported by grant DMS-1407548 from the National Science Foundation, and grant 5R01 EB 001988-21 from the National Institutes of Health.

This research has been conducted using the UK Biobank Resource under application number 24983. We thank all the participants in the study. The primary and processed data used to generate the analyses presented here are available in the UK Biobank access management system (https://amsportal.ukbiobank.ac.uk/) for application 24983, “Generating effective therapeutic hypotheses from genomic and hospital linkage data” (http://www.ukbiobank.ac.uk/wp-content/uploads/2017/06/24983-Dr-Manuel-Rivas.pdf).

All of the computing for this project was performed on the Nero and Sherlock clusters. We would like to thank Stanford University and the Stanford Research Computing Center for providing computational resources and support that contributed to these research results.

